# Visual working memory models of delayed estimation do not generalize to whole-report tasks

**DOI:** 10.1101/2023.03.22.533826

**Authors:** Benjamin Cuthbert, Dominic Standage, Martin Paré, Gunnar Blohm

## Abstract

Whole-report working memory tasks provide a measure of recall for all stimuli in a trial, and afford single-trial analyses that are not possible with single-report delayed estimation tasks. However, most whole-report studies assume that trial stimuli are encoded and reported independently, and do not consider the relationships between stimuli presented and reported within the same trial. Here, we present the results of two independently conducted whole-report experiments. The first dataset was recorded by Adam, Vogel, and Awh, 2017, and required participants to report color and orientation stimuli using a continuous response wheel. We recorded the second dataset, which required participants to report color stimuli using a set of discrete buttons. We find that participants often group their reports by color similarity, contradicting the assumption of independence implicit in most encoding models of working memory. Next, we show that this behavior is consistent across participants and experiments when reporting color but not orientation, two circular variables often assumed to be equivalent. Finally, we implement an alternative to independent encoding where stimuli are encoded as a hierarchical Bayesian ensemble, and show that this model predicts biases that are not present in either dataset. Our results suggest that assumptions made by both independent and hierarchical ensemble encoding models—which were developed in the context of single-report delayed estimation tasks—do not hold for the whole-report task. This failure to generalize highlights the need to consider variations in task structure when inferring fundamental principles of visual working memory.

## Introduction

Many recent models of visual working memory (VWM) were developed in the context of the delayed estimation task (Wilken and Ma, 2004), where participants report the value of a presented stimulus—such as the color of a square or orientation of a bar—after a blank delay period. In this task, report error is measured by taking the difference between presented and reported values, and error distributions over many trials are assumed to reflect the structure of stimulus encodings. Models that reproduce delayed estimation error distributions have therefore been used to support and refute numerous claims about VWM encoding (Zhang and Luck, 2008; Bays and Husain, 2008; T. F. Brady and Alvarez, 2011; van den Berg et al., 2012; Swan and Wyble, 2014; Oberauer, 2017; Nassar, 2018; Schurgin, Wixted, and T. F. Brady, 2020).

Recently, debate over the existence of a fixed storage limit in VWM led Adam and colleagues to introduce a novel delayed estimation variant: the *whole-report task* (Adam, Mance, et al., 2015; Adam, Vogel, and Awh, 2017). Unlike typical delayed estimation tasks—where only one stimulus is reported—each trial of the whole-report task requires participants to report all presented stimuli. This has the advantage of measuring recall of the entire stimulus display, and allowed Adam et al. to investigate item limits at the single-trial level.

The whole-report task is now seeing more widespread use (Peters, Rahm, Czoschke, et al., 2018; Adam and Vogel, 2018; Killebrew et al., 2018; Peters, Rahm, Kaiser, et al., 2019; deBettencourt et al., 2019; Robison and Unsworth, 2019; Utochkin and T. F. Brady, 2020; Hao et al., 2021), and the original publicly-available dataset has been used for model evaluation (Schneegans, Taylor, and Bays, 2020). Critically, these studies treat the whole-report task as an extension of single-report delayed estimation, focusing on error distributions accumulated across trials and reports. Only a few studies report within-trial effects like inter-item interference and bias (Utochkin and T. F. Brady, 2020; Hao et al., 2021; Udale et al., 2021), and none consider the joint distribution of reports made within the same trial.

Within-trial joint distributions are a key affordance of the whole-report task because they allow us to test VWM model assumptions in ways that are not possible with single-report delayed estimation data. For example, many models assume that stimuli are encoded in VWM independently (Zhang and Luck, 2008; Bays and Husain, 2008; van den Berg et al., 2012; Fougnie, Suchow, and Alvarez, 2012; Schneegans, Taylor, and Bays, 2020). If this were the case, a given stimulus encoding would have no effect on other stimuli encoded within the same trial, and the joint distribution of reports would be uniform. Alternative models suggest that stimuli are encoded together in an “ensemble” (Alvarez and Oliva, 2009; T. F. Brady and Alvarez, 2011; Orhan and Jacobs, 2013; Nassar, 2018), which would result in dependencies in within-trial joint distributions.

Here, we investigate previously uncharacterized within-trial behavior in the whole-report task, and find that neither independent nor hierarchical ensemble encoding models can explain the results. We use two whole-report datasets: the original dataset recorded by Adam, Vogel, and Awh, 2017, and data from a whole-report task variant independently conducted in our lab (Cuthbert et al., 2018). In both datasets, we find strong evidence that color reports made in the same trial are not independent. In many cases, participants grouped consecutive within-trial reports by color similarity. In addition, reports made later in the trial—past the canonical capacity limit of 3-4 items—were consistently biased away from colors reported earlier in the trial.

We show that this effect is consistent across participants and set sizes when recalling color stimuli, but is either weaker or absent in task conditions with orientation stimuli. This is surprising, because color and orientation are typically treated as equivalent circular variables (van den Berg et al., 2012; Schneegans, Taylor, and Bays, 2020), and aggregate error distributions for color and orientation stimuli are qualitatively very similar.

Finally, we consider whether hierarchical ensemble encoding models might account for within-trial dependencies in color reports. In these models, dependencies arise due to Bayesian integration over a hierarchical prior (T. F. Brady and Alvarez, 2011; Orhan and Jacobs, 2013). We implement multiple Bayesian ensemble models, and find no evidence for any form of their predicted biases—even after restricting our analyses to specific set sizes or contexts where ensembles might confer a significant storage advantage.

Taken together, our results suggest that whole-report task data cannot be entirely accounted for by VWM models that assume independent encoding, equivalence between circular stimuli, or hierarchical ensemble encoding.

## Methods

We analyzed two whole-report datasets, which we refer to as the *discrete whole-report* and *continuous whole-report* datasets.

### Discrete whole-report dataset

The discrete whole-report dataset was recorded at Queen’s University, and has been previously presented in conference proceedings (Cuthbert et al., 2018).

#### Task and participants

The discrete whole-report task was adapted from Adam, Mance, et al., 2015. 16 healthy adult participants (18-40 years of age; 7 female) completed two experimental sessions each, with 12 blocks of 30 trials completed per session. Participants were required to memorize an array of colored squares (“stimuli”), then report the color of each stimulus after a retention period.

Each trial began with the simultaneous presentation of 2, 3, 4, 6, or 8 stimuli for 500 ms. This was followed by a blank grey screen, presented for 1000 ms, during which participants were required to maintain fixation on the central point. Subjects were then presented with “response matrices,” each comprised of 8 selectable buttons (one for each possible color; see Fig 1). All response matrices were identical within each trial, but button colors were randomized between trials. Response matrices appeared at all locations previously occupied by stimuli, and participants were required to report the color of each stimulus.

**Figure 1.**
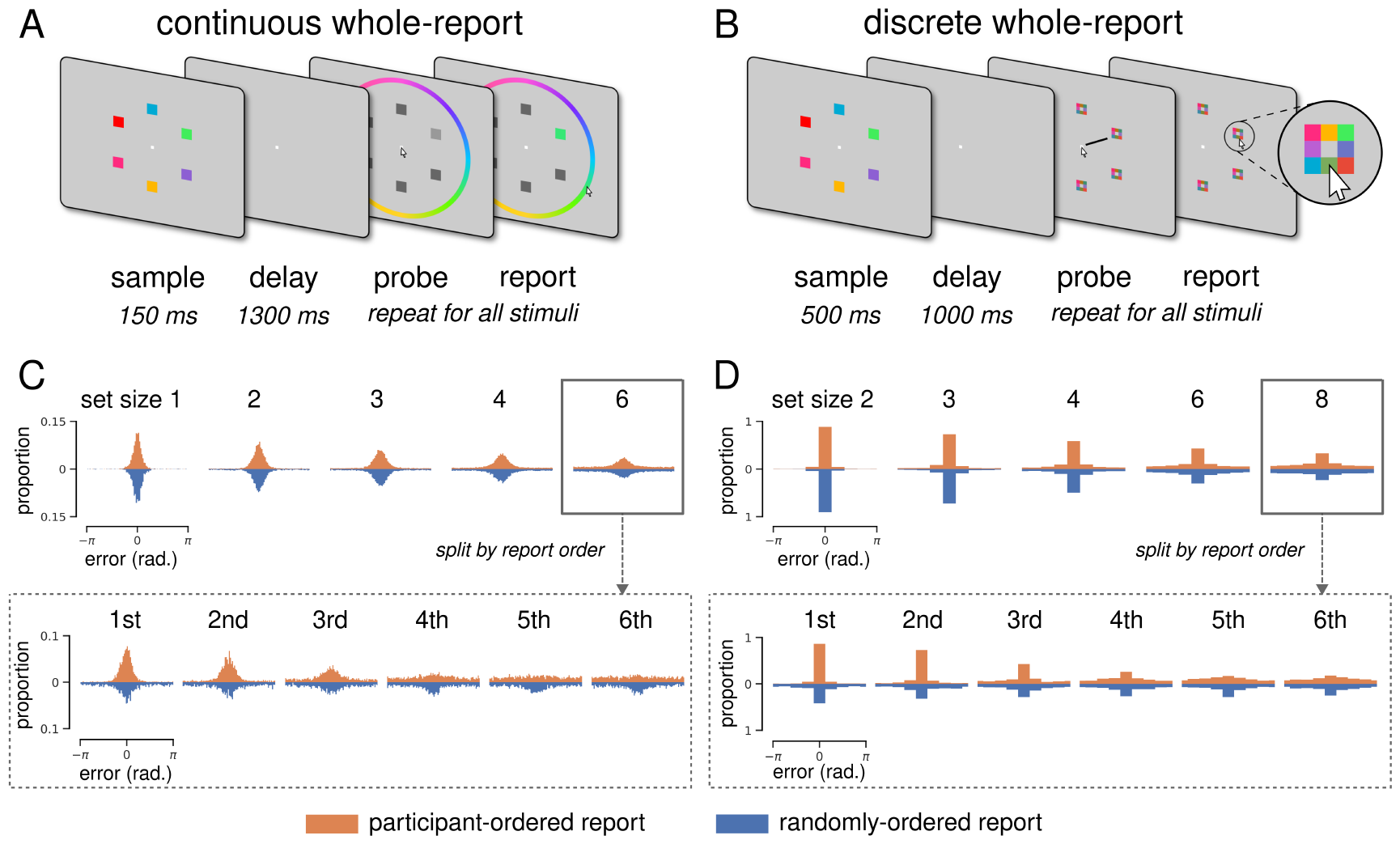
Whole-report task comparison. **A**. Continuous whole-report task. Colored squares were briefly presented, and after a blank delay period the participant reported all colors by clicking a color wheel. Report order was either participant-selected or randomly-generated. Stimuli were sampled from 360 “continuous” color values. **B**. Discrete whole-report task. Same as A, but participants reported by clicking a square in the response array (*inset*). Stimuli were sampled from 8 “discrete” color values. **C**. Continuous error distributions. 360 possible error values divided into 90 bins for visualization. Orange and blue histograms show errors from participant-selected and randomly-generated report conditions, respectively. *Upper*: All report errors, separated by set size. *Lower*: Set size 6 report errors, separated by report order. **D**. Discrete error distributions. Same as C, but bins correspond to all possible error values.

There were two report conditions (one session each): in the *participant-ordered* report condition, participants were instructed to report stimulus colors in order of confidence; in the *randomly-ordered* report condition report order was random, and cued with a black radial line. Color reporting was unspeeded, and each trial concluded once all stimuli had been reported.

#### Experimental setup

Stimuli were generated with MATLAB R2016a (MATLAB, 2016), and presented on a VIEWPixx 3D LCD monitor (VPixx Technologies, Saint-Bruno, QC) using the Psychophysics Toolbox (Kleiner et al., 2007). Compliance during stimulus presentation and the retention period of the task was verified using the EyeLink 1000 Tower Mount (SR Research, Ottawa, ON). The sampling rate was set to 1000 Hz, and eye movement traces of position and velocity were examined for exclusion criteria. Trials were excluded if subjects did not maintain fixation during stimulus presentation (blinks not permitted) and the retention period (blinks permitted). In total, 771 trials were excluded (13.4%).

#### Stimulus generation

Colours were selected from 8 equidistant points on a circle in CIE *L*a*b\** colour space centered at (*L\** = 70, *a\** = 20, *b\** = 38) with a radius of 60. Luminance and chromaticity was calibrated prior to each experimental session using a ColorCAL MKII colorimeter (Cambridge Research Systems Ltd., Rochester, UK), and the coordinates above were selected to ensure that all colors fell within the LCD monitor’s gamut. Stimuli each measured 1.5 x 1.5 ^º^ (visual angle), and were spaced equidistantly around a central fixation point with a 7.5 ^º^ radius.

### Continuous whole-report dataset

The continuous whole-report dataset was recorded by Kirsten Adam and colleagues at the Universities of Chicago and Oregon. The following is a brief summary of their methods, with particular emphasis on the differences between their task and the discrete whole-report task above. For complete experimental details, see Adam, Vogel, and Awh, 2017.

#### Task and participants

Here, we consider data from four task conditions recorded by Adam, Vogel, and Awh, 2017: participant-ordered report with color stimuli (22 participants), randomly-ordered report with color stimuli (17 participants), participant-ordered report with orientation stimuli (23 participants), and randomly-ordered report with orientation stimuli (21 participants). The continuous task used set sizes 1, 2, 3, 4, and 6 (vs 2, 3, 4, 6, and 8 for the discrete task). There were also timing differences; continuous task stimuli were presented for 150 ms (vs 500 ms), and the delay period lasted 1300 ms (vs 1000 ms).

#### Discrete vs continuous stimuli

While color stimuli for both tasks were sampled from equidistant points around circles in CIE *L*a*b\** color space, the discrete task only used 8 distinct colors while the continuous task used 360. The area of color space used was also different—the continuous task used a circle centered at (*L\** = 54, *a\** = 18, *b\** = -8)—and monitors were not calibrated before the continuous experiment, so the displayed colors cannot be considered perceptually equivalent to discrete task stimuli. The continuous task also included conditions with orientation stimuli, which were sampled from 360 equally-spaced angles.

#### Discrete vs continuous report method

As illustrated in Figure 1, the two tasks employed different report methods. In the discrete task, 8 possible report colors were presented in the form of clickable buttons at the stimulus location being probed. In the continuous task, 360 possible report colors were presented in the form of a clickable color wheel surrounding the entire display. Similarly, the continuous task with orientation stimuli required participants to click on a surrounding (uncolored) wheel to report remembered orientation.

### Analyses

#### Kullback-Leibler (KL) divergence estimation

To quantify within-trial dependence, we estimated the KL divergence between within-trial relative distance distributions and a circular uniform distribution. KL divergence is a measure of the difference beteen two distributions that involves computing their relative entropy (in bits). A low KL divergence between a discrete distribution *P* and a nominal uniform distribution *Q* is evidence that *P* has high entropy—and is close to uniformly distributed. A high KL divergence provides evidence that *P* is not uniformly distributed. This approach is useful because unlike correlation, it does not assume a linear relationship between variables, and can therefore capture higher-order dependencies such as those evident in Fig. 2.

**Figure 2.**
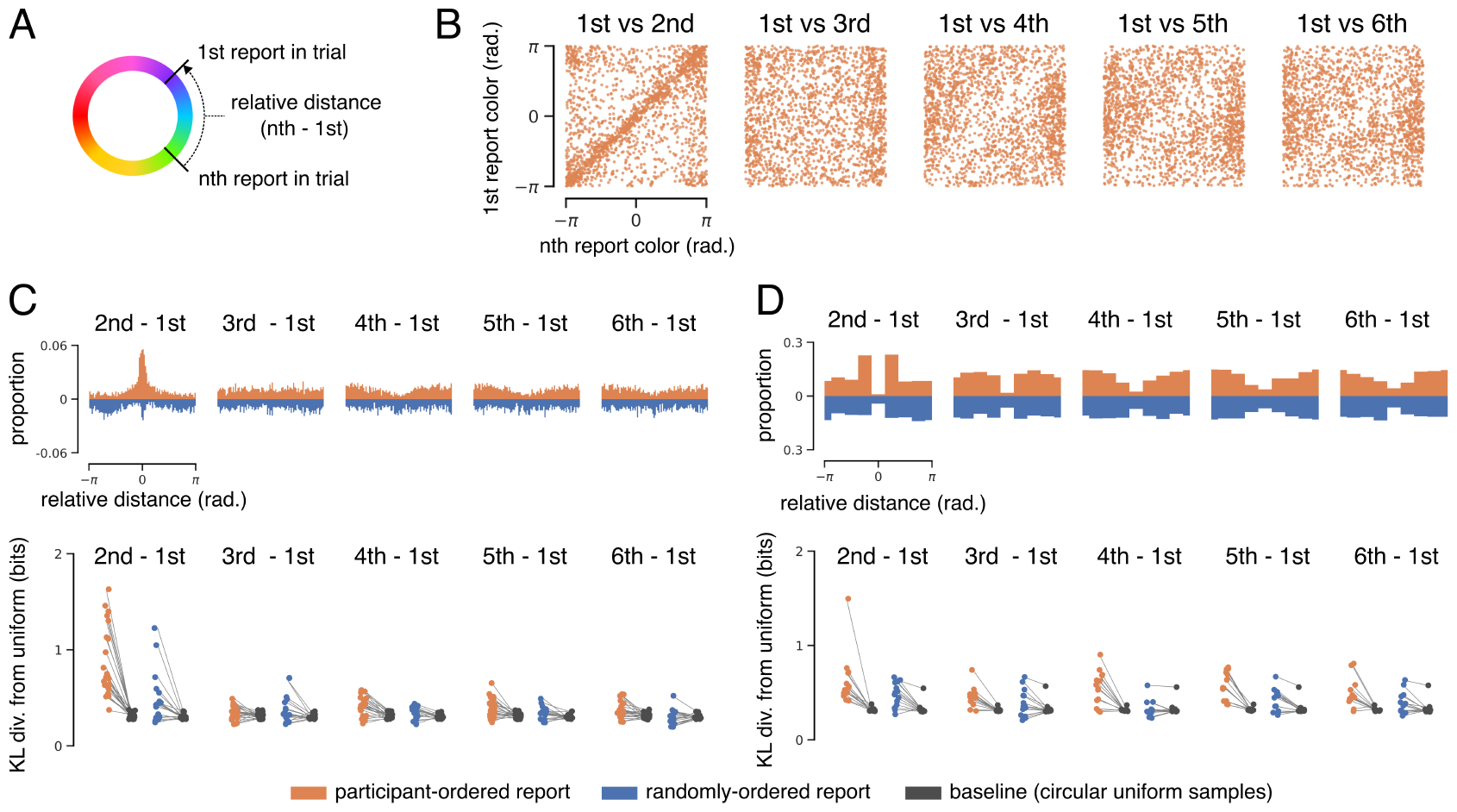
Dependence of within-trial color reports. **A**. Relative distance calculation, which was identical for both tasks. **B**. Example joint distribution of reported stimulus values. Data from the participant-ordered continuous whole-report task (set size 6; aggregated across all participants). Each panel plots the 1st color reported on each trial against a later report in the same trial. **C**. *Upper:* Distribution of relative distances for the continuous task (set size 6). *Lower:* Median bootstrapped estimates of the KL divergence between each participant’s relative distance distribution and size-matched samples from a uniform distribution. **D**. Same as C, but for the discrete whole-report task (set size 6).

For each participant, report condition, and pair of within-trial reports, we performed 1000 boot-strapped KL divergence estimates between the empirical distribution *P* and a nominal circular uniform distribution *Q*:

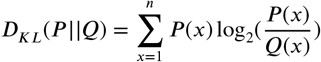

where *n* is the number of equal bins used to define the probability space (*n* = 36 for the continuous task and *n* = 8 for the discrete task).

To generate a baseline estimate, we repeated this process but replaced each empirical distribution *P* with a size-matched sample from a circular uniform distribution. Median estimates for all participants are shown alongside their respective baseline median estimates in Fig. 2C and D.

### Hierarchical encoding models

#### Two-level Hierarchical Bayesian Model (HBM)

This model is adapted from the simplest model introduced by T. F. Brady and Alvarez, 2011. In the HBM implemented here, stimuli are probabilistically represented in working memory at two levels: (1) individual stimulus color estimates and (2) a joint “ensemble” of all stimuli.

Here, both color and orientation are considered circular stimulus spaces, so we substitute the original Gaussian distributions with the von Mises (circular normal) distribution:

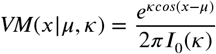

where *I*_0_(⋅) is the modified Bessel function of order 0, *μ* is the mean circular direction, and *κ* is the concentration parameter (analagous to 1/*σ*^2^).

The HBM assumes that on each trial, *N* stimuli 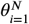 are sampled from the same “ensemble” von Mises distribution:

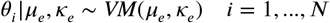

where *μ*_*e*_ and *κ*_*e*_ are the ensemble mean and precision with the following priors:

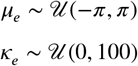

The HBM observer only has access to noisy von Mises observations of 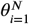:

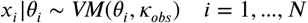

where *κ*_*obs*_ is the precision of noisy observations 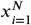 centered on 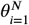.

*κ*_*obs*_ is therefore the only free parameter in this model. We estimated plausible values of *κ*_*obs*_ by fitting a von Mises function—via maximum likelihood estimation—to error distributions from set size 1 of the continuous whole-report task (see Results; Fig. 3B). To generate a range of model predictions, all simulations were conducted in triplicate using *κ*_*obs*_ = 5, 10, 20.

**Figure 3.**
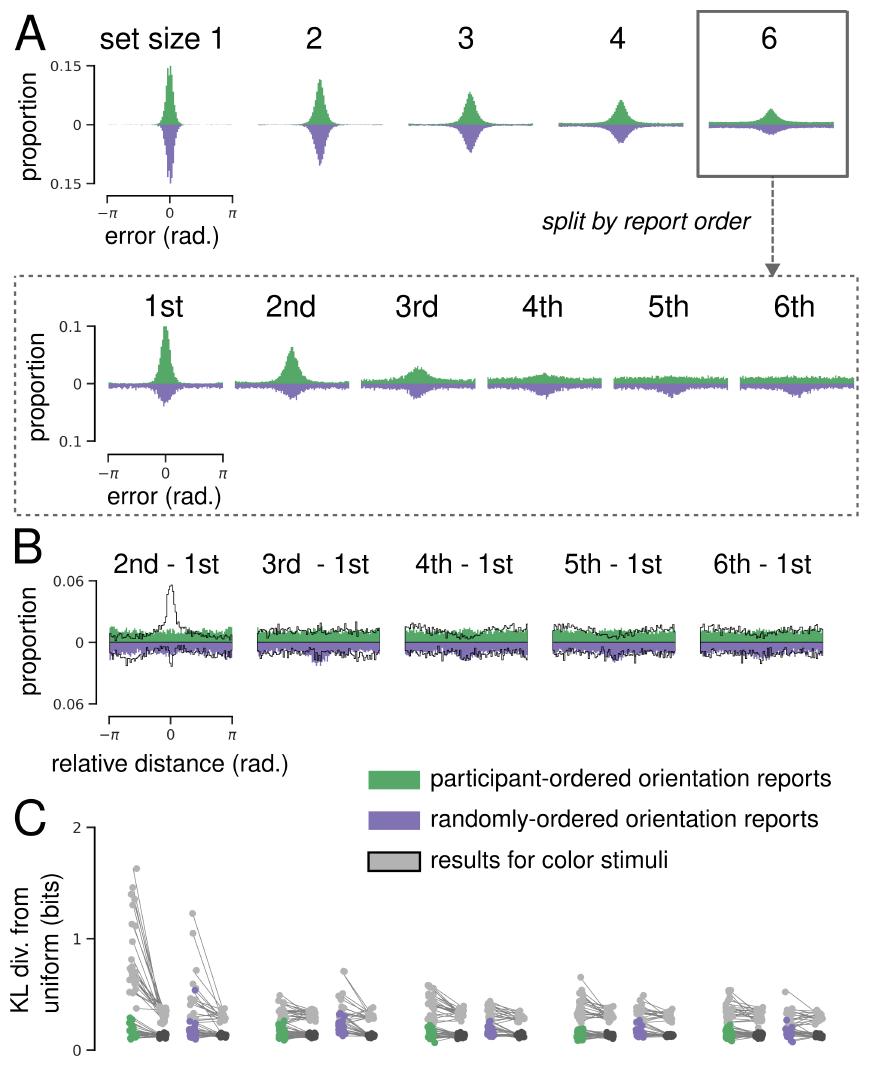
Continuous whole-report results for orientation stimuli. **A**. Orientation error distributions. 360 possible error values divided into 90 bins for visualization. Green and purple histograms show errors from participant-selected and randomly-generated report conditions, respectively. *Upper*: All report errors, separated by set size. *Lower*: Set size 6 report errors, separated by report order. **B**. Distribution of relative distances for orientation reports (set size 6). Color results overlayed in black for comparison. **C**. Median bootstrapped estimates of the KL divergence between each participant’s relative distance distribution and size-matched samples from a uniform distribution. Color results overlayed in grey for comparison.

#### Bayesian Finite Mixture Model (BFMM)

We also implemented a more complex generalization of the HBM known as a Bayesian finite mixture model (Orhan and Jacobs, 2013). In addition to individual- and ensemble-level representations, the BFMM also includes an intermediate level where stimuli are generated from a set of *K* components, such that similar stimulus values are assumed to be generated from the same component. Performing inference with the BFMM can be thought of as finding a posterior distribution over all possible assignments of *N* stimuli to *K* clusters.

Formally, each stimulus *θ*_*i*_ is assumed to be generated by one of *K* weighted von Mises components. The components are assumed to be generated by the following process:

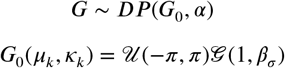

where G is a discrete distribution over the parameters of the *K* components. *G*_0_ is the prior distribution over the two-dimensional parameter space for *μ*_*k*_ and *κ*_*k*_, the mean and precision of component *k*. The *β*_*σ*_ variable for the gamma prior is given a 𝒢(1, 1) hyperprior.

G can be expressed as a weighted sum of *K* “atoms” or points *p* in the two-dimensional parameter space *G*_0_:

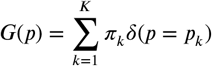

where *π*_*k*_ is the weighting of component *k* and *δ* is the Dirac delta function. The component weights *π* are drawn from a symmetric Dirichlet prior with concentration parameters *α*/*K*:

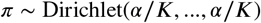

The assumed stimulus-generating and observation processes are then:

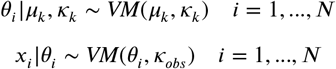

where 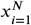 are noisy von Mises observations centered on true stimulus values 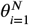 with precision *κ*_*obs*_.

The BFMM has two free parameters: *κ*_*obs*_ indicating the observation precision, and the Dirichlet clustering parameter *α*. As above, simulations were performed in triplicate with *σ*_*obs*_ = 5, 10, 20. The clustering parameter *α* was fixed at 1 for all simulations, roughly corresponding to a uniform distribution over component weights (Orhan and Jacobs, 2013).

#### Model simulations

Model simulations were performed in the same way for both the HBM and the BFMM. To simulate a whole-report trial at set size *N*, a noisy estimate was first generated for each stimulus by sampling from a von Mises distribution with precision *κ*_*obs*_. The models do not have access to the true values of *θ*, but instead perform inference based on these noisy observations.

Inference and model specification were implemented using the probabilistic Python package PyMC3 (Salvatier, Wiecki, and Fonnesbeck, 2016). Model posteriors were sampled using a Hamiltonian Monte Carlo algorithm known as the No-U-Turn Sampler (Hoffman and Gelman, 2014). For each trial, 5000 samples were drawn (after 2000 tuning steps).

To simulate reporting, the mode of each stimulus posterior was used as a point estimate (taking the mean did not change our results), and point estimates were rounded to one of either 360 or 8 possible values for the continuous and discrete task, respectively.

### Data and code availability

All Python code required to reproduce data processing, analysis, modeling, and visualization is available at this project’s [GitHub repository](https://github.com/bncuthbert/whole-report-wm). The repository also contains both datasets in their original raw `.mat` format, as well as pickled binary versions (stored in pandas Dataframes) suitable for analysis with Python. Installation and usage instructions are provided, and all code is thoroughly documented.

## Results

We analyzed behavioural data from two independent whole-report experiments to characterize within-trial report behavior and evaluate the applicability of encoding models developed in the context of single-report tasks.

These whole-report experiments used two subtly different tasks (Fig. 1A-B; Methods), so we began by replicating the continuous whole-report results of Adam, Vogel, and Awh, 2017 with discrete whole-report data. We found that cumulative error distributions were qualitatively similar despite differences in experimental setup, stimuli, timing, and report method.

We then analyzed within-trial behavior, and found that participants tended to group consecutive reports by color similarity—reports made later in a trial were often biased away from early-trial reports. This pattern of dependence was consistent across datasets, participants, and set sizes when reporting color, but not when reporting orientation.

Finally, we implemented two hierarchical Bayesian ensemble encoding models to determine whether color report dependecies could be explained by the joint encoding of within-trial stimuli. We simulated both whole-report tasks using both models, and found no empirical evidence for the biases that either model predicted.

### Continuous and discrete whole-report error distributions are qualitatively similar

In order to verify that differences in task timing, stimuli, and report method between the continuous and discrete tasks did not impact participant behavior, we replicated the aggregate results of Adam, Vogel, and Awh, 2017 with the discrete whole-report dataset. When comparing data between tasks, we considered both report order conditions (participant- and randomly-ordered), and restricted continuous task data to conditions using color stimuli (the discrete experiment did not include orientation stimuli).

Error distributions from both tasks—collapsed across trials and participants—are shown in Fig. 1. A classic delayed estimation result is that recall performance decreases as a function of set size, and Adam, Vogel, and Awh, 2017 found that continuous whole-report error distributions follow the same trend. Error distributions are less precise at higher set sizes (Fig. 1C; upper) in both the participant- and randomly-ordered report conditions.

Discrete whole-report error distributions are qualitatively similar (Fig. 1D; upper). Following Adam, Vogel, and Awh, 2017, we quantified this effect using mean resultant vector length (MRVL), a measure of circular dispersion. MRVL decreases as a function of set size in both report conditions, and repeated-measures analyses of variance (ANOVAs) show a significant effect of set size on MRVL in both the participant-ordered condition (F(4,52) = 339.66, *p* = 1.40 × 10^−36^) and randomly-ordered condition (F(4,52) = 268.22, *p* = 5.00 × 10^−34^).

Whole-report error distributions can be further separated by the order in which stimuli were reported within each trial. This is illustrated for set size 6 in the bottom panels of Fig. 1C and D (see Supplementary Fig. 8 for all set sizes). Adam, Vogel, and Awh, 2017 found that when report order is freely selected by participants, MRVL decreases as a function of report number (reproduced in Supplementary Fig. 9A).

Discrete whole-report error distributions remain similar to continuous distributions when separated by report order. In the participant-ordered condition, MRVL decreases as a function of report number (Supplementary Fig. 9B), and repeated-measures ANOVA showed a significant effect of report order for all set sizes. Notably, in this condition discrete whole-report participants were instructed to report stimuli “in order of confidence,” and participants in the continuous task appear to have done the same despite receiving no instruction.

### Within-trial color reports are not independent

As discussed above, the whole-report task allows us to analyze the joint distribution of stimulus reports within a trial. Several models of delayed estimation make the assumption that stimuli are encoded independently—if this is the case, the joint distribution of within-trial reports should be uniform.

Fig. 2B shows example within-trial joint distributions for set size 6 of the participant-ordered continuous whole-report task. Each panel corresponds to the joint distribution between the 1^*st*^ and the *n*^*th*^ color reported within the same trial. ^1^ Two dependencies are apparent: participants tended to report similar colors for the 1^*st*^ and 2^*nd*^ reports in a trial, and they tended to avoid reporting colors similar to the 1^*st*^ with later reports. This is not unique to the 1^*st*^ report—consecutive reports are similar regardless of when in the trial they occurred.

Another way to visualize this is to compute the relative angular distance between two colors reported in the same trial (Fig. 2A). The upper panel of Figure 2C shows relative distance distributions computed for the continuous whole-report task at set size 6. The pattern of dependencies described above is evident for the participant-ordered condition. Deviations from uniformity are also evident for the randomly-ordered report condition, despite the fact that the corresponding distributions of *presented* values were uniform. Critically, this effect was not driven by a few particpants making idiosyncratric reports, but was consistent across all participants (Supplementary Fig. 10) and set sizes (Supplementary Fig. 11).

Deviations from uniformity are evident for the discrete whole-report task, and patterns of dependence were qualitatively similar to those evident in the continuous task data, with one key difference—in both conditions of the discrete task, participants rarely repeated their 1^*st*^ report later in the trial. This suggests that participants were not simply reporting the same color repeatedly, but explicitly avoiding previous colors while consecutively reporting similar colors.

We quantified dependence between within-trial reports using Kullback-Leibler (KL) divergence, a measure of the statistical distance between two distributions. For each participant, report condition, and pair of reports, we estimated the KL divergence between the relative distance distribution and a circular uniform distribution. We compared these KL divergence estimates to a baseline divergence obtained by repeating this process using a size-matched circular uniform sample. This process was bootstrapped to obtain an estimate of uncertainty (see Methods for details), and median KL divergence estimates for each participant are shown alongside corresponding baseline estimates in the lower panels of Fig. 2C and D. This analysis confirmed that distributions of within-trial relative distance often deviated from uniformity for all participants of both tasks. Divergences tended to be higher in the participant-ordered condition than in the randomly-ordered condition, and were highest for consecutive reports.

Repeated-measures ANOVAs using estimate type (ie. data vs baseline) and report pair as within-subject factors confirmed a significant effect of estimate type on KL divergence in both the particpant-ordered (F(1,21) = 70.49, *p* = 3.76×10^−8^) and randomly-ordered (F(1,16) = 10.55, *p* = 0.005) conditions of the continuous task. The same analysis confirmed a significant effect in both the particpant-ordered (F(1,13) = 55.13, *p* = 5.0 × 10^−6^) and randomly-ordered (F(1,13) = 32.74, *p* = 7.0 × 10^−5^) conditions of the discrete task. These analyses also showed significant effects of report order, as well as an interaction between report order and estimate type.

### Within-trial orientation reports do not show clear dependence

Many models of delayed-estimation tacitly assume that different types of “circular” stimuli—such as colored squares and orientated barscare equivalently encoded into working memory. This is supported by qualitative similarities in error distributions in single-report delayed estimation tasks (Zhang and Luck, 2008; van den Berg et al., 2012; Bays, Catalao, and Husain, 2009). Adam, Vogel, and Awh, 2017 found that aggregate error distributions for orientation and color stimuli were qualitatively similar in the whole-report task. Error distributions for reported orientations—collapsed across reports and participants—are less precise at higher set sizes (Fig. 3A; upper) in both the participant- and randomly-ordered report conditions. They also found that when report order was freely selected by participants, MRVL decreased as a function of report number—again mirroring results for color stimuli.

Despite this similarity at the level of aggregate errors, we found marked differences between color and orientation when analyzing within-trial reports (Fig. 3B and C). Relative distance distributions for orientation do not exhibit clear dependence between consecutive reports in the participant-ordered condition, nor did participants avoid repeating early reports later in the trial. As above, we quantified dependence between within-trial orientation reports using Kullback-Leibler (KL) divergence, a measure of the difference between two distributions. Median KL divergence estimates for each participant are shown alongside corresponding baseline estimates in Fig. 3C (results for color reports are overlayed in gray for comparison). ^2^ In contrast to results for color (Fig. 2), this analysis revealed that for most participants, deviations from uniform were either greatly reduced or absent.

We performed repeated-measures ANOVAs using estimate type (ie. data vs baseline) and report pair as within-subject factors. This analysis found a significant effect of estimate type on KL divergence in both the particpant-ordered (F(1,29) = 39.10, *p* = 0.01 × 10^−4^) and randomly-ordered (F(1,18) = 34.12, *p* = 1.6 × 10^−5^) conditions, but no significant effects of report order or interaction between report order and estimate type. Although this suggests some dependence between orientation reports, this effect was greatly reduced in comparison to results for color.

### Hierarchical ensemble encoding predicts biases not present in whole-report data

As described above, we found clear evidence that within-trial color reports are not independent in the whole-report task. Participants appear to group stimuli by color when reporting, raising the possibility that they are encoding higher-order statistics of color displays.

This is a proposal made by “ensemble” encoding models of VWM, where visual scenes are hierarchically encoded at multiple levels of abstraction and individual stimulus representations are marginally dependent due to shared ensemble statistics (T. F. Brady, Konkle, and Alvarez, 2009; T. F. Brady and Alvarez, 2011; Orhan and Jacobs, 2013). This introduces potentially unwanted biases, but could reduce variance in stimulus estimates (Orhan and Jacobs, 2013) and improve encoding efficiency (Nassar, 2018).

We implemented two hierarchical encoding models to simulate the whole-report task and test this possibility. The first is a simple adaptation of T. F. Brady and Alvarez, 2011: a two-level hierarchical Bayesian model (HBM) that encodes both individual stimuli and the ensemble statistics—mean and precision—of all stimuli in the display. This model makes the simple, testable prediction that color reports will be biased toward the mean color of the display.

To address the possibility that participants may be encoding multiple groups of similar colors, we also adapted a hierarchical encoding model known as a Bayesian finite mixture model (BFMM) (Orhan and Jacobs, 2013). The BFMM has another level of abstraction where stimuli are grouped by similarity, and the statistics of each group are also encoded. This model predicts that color reports will be biased toward the mean of the group to which they belong.

#### Hierarchical Bayesian model (HBM)

The biases produced by a HBM with two levels of representation are illustrated in Fig. 4A. In the HBM, an observer assumes that all stimuli in a trial are samples from the same underlying ensemble distribution, and this internal model effectively forms a prior over the values of all stimuli. When integrating over the uncertainty in individual stimulus encodings and this prior, the observer produces posterior estimates that are biased toward the ensemble mean.

**Figure 4.**
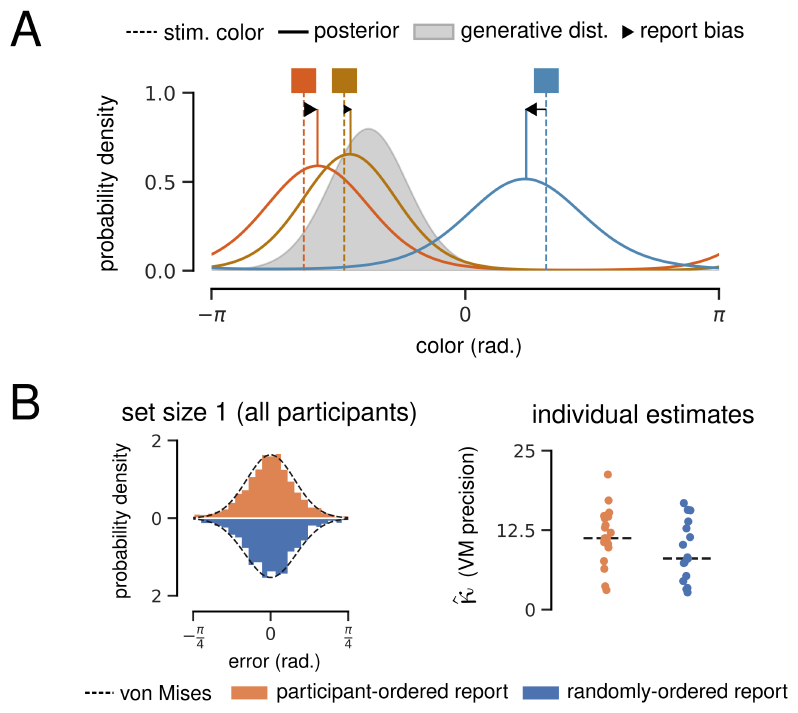
Hierarchical Bayesian Model (HBM). **A**. Illustration of the HBM encoding 3 color stimuli (vertical dashed lines). The model infers that all colors are drawn from the same ensemble distribution (grey). Solid lines show posterior estimates for each stimulus value, and black arrows illustrate the resulting bias toward the ensemble mean. **B**. Justification for free parameter *κ*_obs_, the precision of noisy observations. *Left:* Error distributions for color reports at set size 1. 360 error values divided into 90 bins for visualization. Maximum-likelihood von Mises distribution fit overlayed. *Right:* Median bootstrapped estimate of the precision of a von Mises fit to set size 1 report errors for each participant.

The HBM implemented here assumes that individual stimulus and ensemble encodings take the form of a von Mises—or “circular normal”—distribution, each parameterized by a mean (*μ*) and precision (*κ*). These parameters are all inferred from presented stimuli via Bayesian inference (see Methods for full equations and priors). The model also assumes that stimulus values are only available as a noisy observation sampled from a von Mises distribution with precision *κ*_*obs*_ centered on the true value. *κ*_*obs*_ is the HBM’s only free parameter, and is assumed to represent the combined effect of sensory and memory noise (Orhan and Jacobs, 2013). Intuitively, lower values of *κ*_*obs*_ result in noisier individual representations and a greater bias toward the ensemble mean.

To estimate reasonable values for *κ*_*obs*_, we took advantage of continuous whole-report error distributions at set size 1, where there are theoretically no higher-order statistics to encode. We fit a von Mises distribution to each participant’s report errors at set size 1, giving us a range of plausible *κ*_*obs*_ values (Fig. 4B). For all HBM and BFMM results presented here, simulations were conducted in triplicate using *κ*_*obs*_ = 5, 10, 20.

Here, we focus on simulation results from set size 3, reasoning that the HBM would tend to infer a more precise generative distribution—and therefore predict stronger biases—with fewer stimuli. HBM simulations of the continous and discrete whole-report tasks are presented in Fig. 5A and C, respectively. Report biases can be visualized by comparing two angles: (1) the angular distance between a presented color and the trial ensemble mean and (2) the error between presented and reported colors. A correlation between the signs of these two angles provides evidence that reported colors are biased toward the ensemble mean.

**Figure 5.**
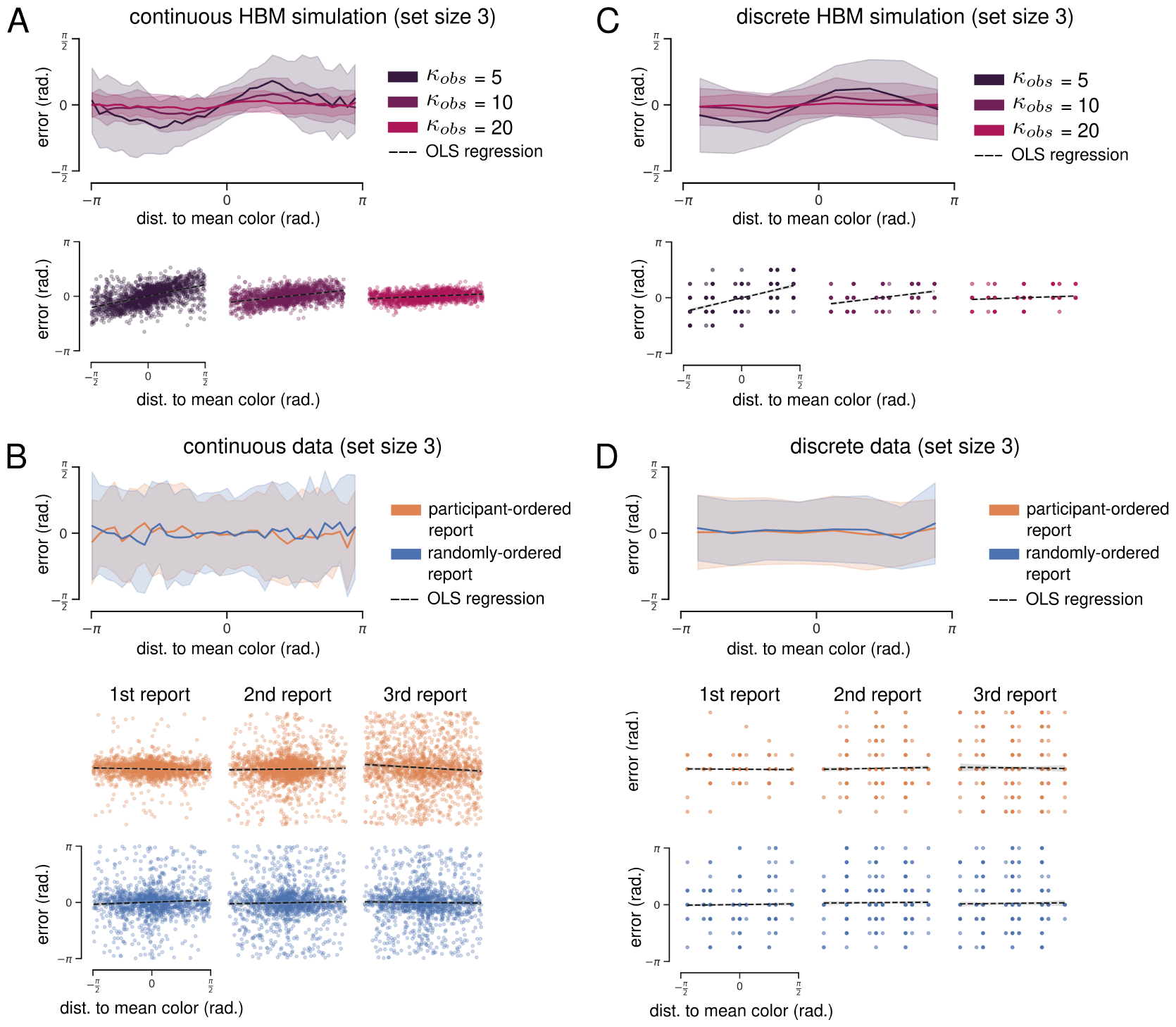
Hierarchical Bayesian model (HBM) simulation results for set size 3. **A**. *Upper:* Biases predicted by HBM simulations of the continuous task. Mean report error plotted as a function of the reported color’s distance to the ensemble mean. 360 error values are divided into 90 bins for visualization, and shaded area shows standard deviation. *Lower:* OLS regression of simulated report error on distance to ensemble mean (dashed black lines). **B**. *Upper:* Empirical biases for the continuous task at set size 3 (collapsed across all reports). *Lower:* OLS regression of empirical report error on distance to ensemble mean. Each column shows results for a different report number. **C**. Same as panel A, but for simulations of the discrete task. **D**. Same as panel B, but for empirical results of the discrete task.

As expected, HBM simulations of both tasks demonstrate a clear bias toward the ensemble mean. Bias magnitude was a function of the distance to the mean color—the farther a stimulus was from the ensemble mean, the greater the reported bias toward the mean. Bias magnitude was also dependent on the choice of *κ*_*obs*_—the lowest *κ*_*obs*_ value resulted in the greatest bias and vice versa. We quantified bias using ordinary least squares (OLS) regression of report error on the distance to ensemble mean. This analysis was restricted to include only mean distances between 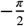 and 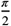 where the bias function was approximately linear (see Methods for details). Regression was done at two levels: (1) all simulated errors for a given set size, collapsed across reports; and (2) simulated errors separated by report number. ^3^

OLS regression was significant for all *κ*_*obs*_ in continuous whole-report HBM simulations at set size 3 (collapsed across reports). The largest estimated effect was for *κ*_*obs*_ = 5 (*R*^2^ = 0.278, mean dist. coefficient = 0.428, *p* = 2.97 × 10^−124^), and the smallest estimated effect for *κ*_*obs*_ = 20 (*R*^2^ = 0.053, mean dist. coefficient = 0.078, *p* = 3.63 × 10^−63^. OLS regressions were significant when simulated reports were separated by report order.

OLS regression was also significant for all *κ*_*obs*_ in discrete whole-report HBM simulations at set size 3 (collapsed across reports). The largest estimated effect was for *κ*_*obs*_ = 5 (*R*^2^ = 0.252, mean dist. coefficient = 0.454, *p* = 4.03 × 10^−104^), and the smallest estimated effect for *κ*_*obs*_ = 20 (*R*^2^ = 0.024, mean dist. coefficient = 0.053, *p* = 2.73 × 10^−10^). OLS regressions were significant when simulated reports were separated by report order.

Empirical bias functions for the continuous and discrete whole-report tasks are shown in Fig. 5B and D, respectively. Empirical report errors were more broadly distributed (less accurate) than simulated results, but report error was comparable when only the first report in each trial was considered (Fig. 5B and D; first column).

We repeated the regression analysis using experimental data from both report conditions of both tasks at set size 3. Regression was significant for the participant-ordered continuous data (collapsed across reports), but the estimated mean distance coefficient was negative, indicating a bias *away* from the ensemble mean (*R*^2^ = 0.001, mean dist. coefficient = −0.050, *p* = 0.004). We also found significant negative biases for the 1^*st*^ and 3^*rd*^ reports when the data were separated by report order. There were no significant regression results for the randomly-ordered condition of the continuous whole-report task at set size 3, or for either condition of the discrete whole-report task.

We also simulated both tasks and report conditions at set size 6, and repeated the OLS regression analysis for both simulated and experimental reports. In comparison to set size 3 simulations, the HBM tended to infer less precise generative distributions, resulting in smaller predicted biases. These biases were still significant for all continuous task simulations, but were not significant for the majority of discrete task simulations—likely because the predicted biases were smaller than the discrete stimulus increment ( ^*π*^ ) and were lost when estimates were rounded for report.

There were several significant regression results for experimental data from both tasks at set size 6. Similar to set size 3, these regressions all estimated small negative mean distance coefficients. Finally, we repeated all analyses above with experimental data from the continuous task with orientation stimuli (Supplementary Fig. 12). We found no significant regressions with positive coefficients in either report condition at set size 3 or set size 6. In summary, we found no empirical evidence for a report bias toward within-trial mean stimulus values.

#### Bayesian finite mixture model (BFMM)

The HBM presented above assumes that all stimuli are generated from a single distribution. The limitations of this approach are clear when considering the HBM illustration in Fig. 4: although encoding orange and red together is intuitive, it is not obvious that the blue stimulus should be included. It is possible that participants cluster similar stimuli together, and encode ensemble statistics of these clusters in addition to statistics of the entire display (T. F. Brady and Alvarez, 2011; Orhan and Jacobs, 2013).

To address this, we adapted a flexible generalization of Brady and Alvarez’s HBM called a Bayesian finite mixture model (BFMM) (Orhan and Jacobs, 2013). The BFMM assumes that each stimulus value is generated from one of *K* weighted von Mises components (Fig. 6A shows an example where *K* = 2). ^4^ Component weights and parameters are determined via Bayesian inference, so sampling from the posterior of the BFMM can be thought of as performing data-driven clustering (see Methods for full model and priors).

**Figure 6.**
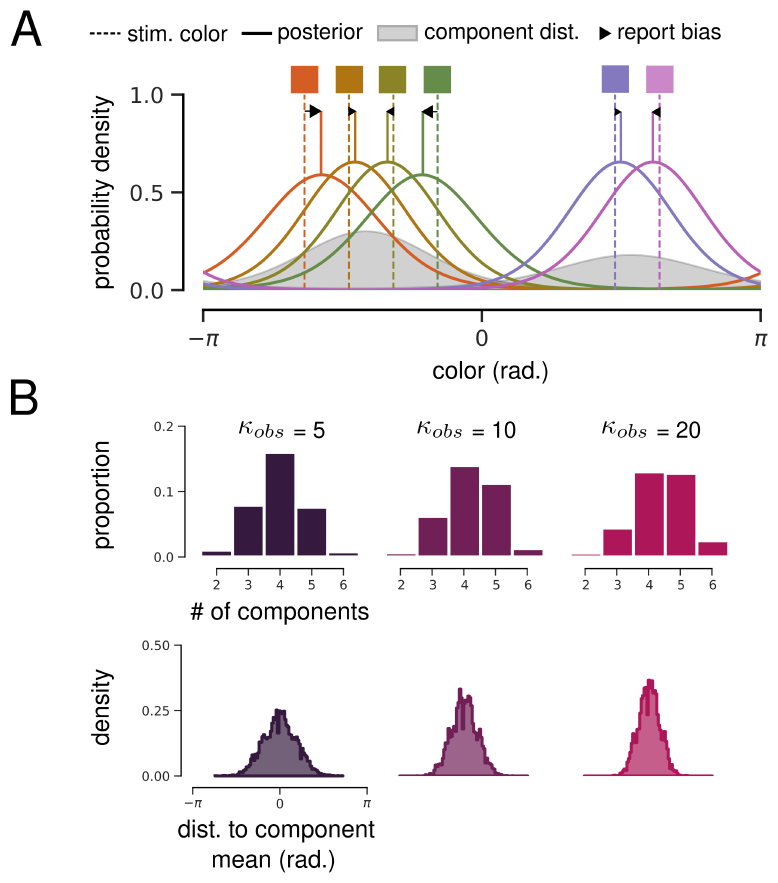
Bayesian finite mixture model (BFMM). **A**. Illustration of a BFMM with *K* = 2 components encoding 6 color stimuli (vertical dashed lines). The model infers that colors were generated from a mixture of 2 von Mises components (grey). Solid lines show posterior estimates for each stimulus value, and black arrows illustrate resulting biases toward component means. **B**. Impact of *κ*_*obs*_ on clustering in a BFMM with *K* = 6 components. *Upper:* Distribution of the number of components that the BFMM infers generated colors on each trial of the simulated continuous whole-report task at set size 6. Higher values of *κ*_*obs*_ cause the BFMM to assume a higher number of more precise components. *Lower:* Distribution of distance to inferred component mean for all simulated reports.

*K* (the maximum number of components) was fixed to equal the simulated set size, and the same values of *κ*_*obs*_ (the precision of noisy observations) were used. Here, *κ*_*obs*_ has a subtly different impact on simulation results than in the HBM (Fig. 6B). At lower values of *κ*_*obs*_, the model tends to infer that stimuli were generated from a small number of low-precision components, whereas at higher values of *κ*_*obs*_, the model tends to infer a greater number of higher-precision components (up to a maximum of *K*).

BFMM simulations were very similar to HBM simulations described above (Fig. 7). To visualize bias, we plot report error as a function of the distance to the mean of the component that the model assumes is responsible for generating a given stimulus. As expected, simulated reports for both the continuous and discrete tasks demonstrate a clear bias toward their respective component means. Similar to the HBM simulations above, bias magnitude was greater for larger distances to the component mean, and was greater for smaller values of *κ*_*obs*_. Overall, BFMM report biases were greater than HBM biases.

**Figure 7.**
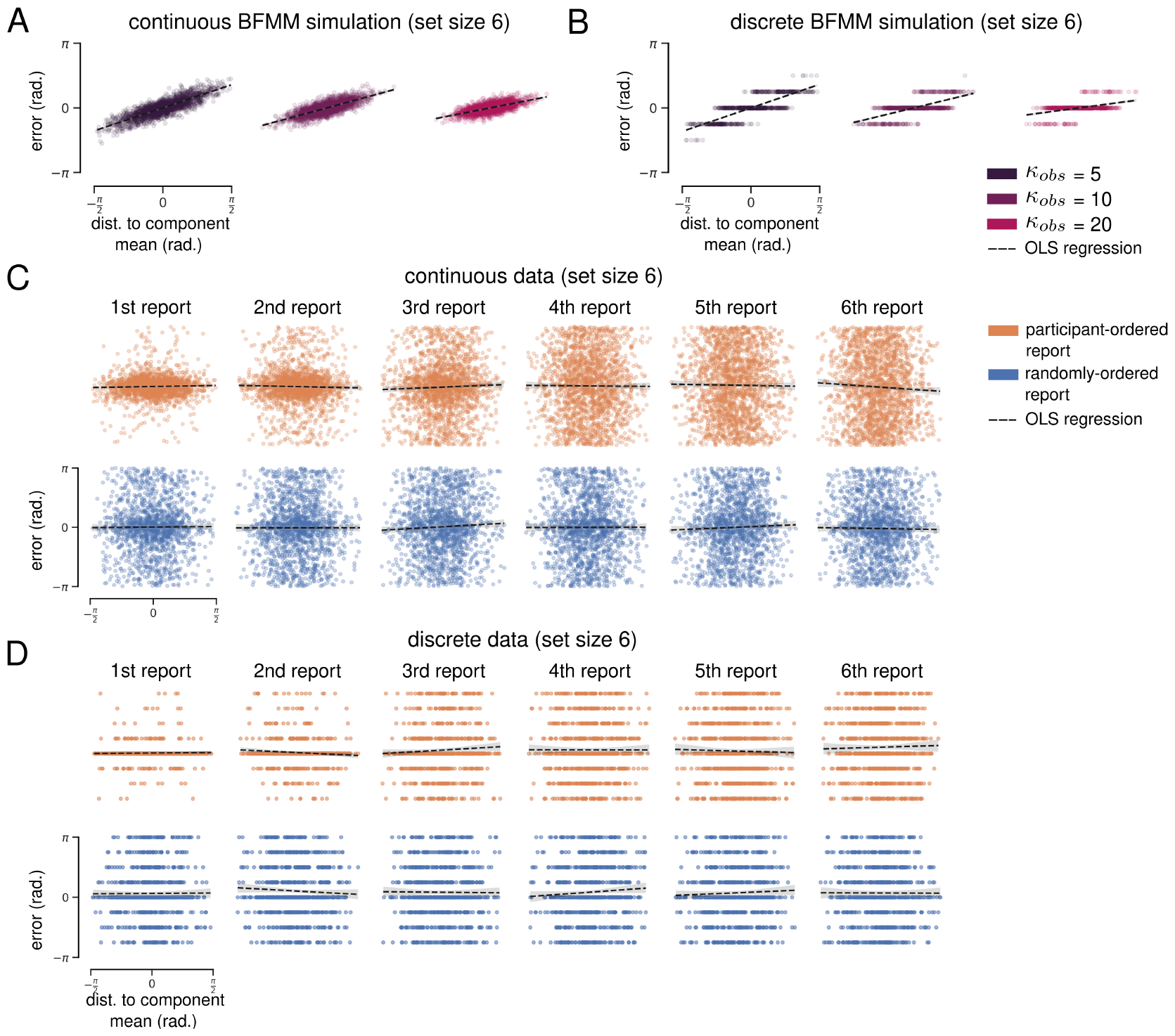
Bayesian finite mixture model (BFMM) simulation results for set size 6. **A**. OLS regression of simulated continuous report error on distance to component mean (dashed black lines). **B**. Same as panel A, but for discrete task simulations. **C**. OLS regression of empirical report error on distance to component mean. Each column shows results for a different report number. **D**. Same as C, but for empricial results of the discrete task.

To quantify bias, we used ordinary least squares (OLS) regression of report error on the distance to component mean. As above, we restricted this analysis to include only mean distances between 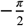 and 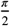 (in practice, this included nearly all reports; see Fig. 6B).

OLS regression was significant for all *κ*_*obs*_ in continuous whole-report BFMM simulations at set size 6 (collapsed across reports). The largest estimated effect was for *κ*_*obs*_ = 5 (*R*^2^ = 0.746, mean dist. coefficient = 0.725, *p <* 1.0 × 10^−300^), and the smallest estimated effect for *κ*_*obs*_ = 20 (*R*^2^ = 0.465, mean dist. coefficient = 0.429, *p <* 1.0 × 10^−300^). OLS regressions were also significant when simulated reports were separated by report order.

OLS regression was also significant for all *κ*_*obs*_ in discrete whole-report BFMM simulations at set size 6 (collapsed across reports). The largest estimated effect was for *κ*_*obs*_ = 5 (*R*^2^ = 0.644, mean dist. coefficient = 0.722, *p <* 1.0 × 10^−300^), and the smallest estimated effect for *κ*_*obs*_ = 20 (*R*^2^ = 0.207, mean dist. coefficient = 0.282, *p* = 1.41 × 10^−247^). OLS regressions were also significant when simulated reports were separated by report order.

We repeated the OLS regressions using empirical data from both report conditions of both tasks at set size 6. Regressions were not significant for either report condition of either task when data were collapsed across reports.

When separated by report order, three regressions reached significance for the continuous whole-report task: the 3^*rd*^ and 6^*th*^ report in the participant-ordered condition and the 3^*rd*^ report of the randomly-ordered condition. These regressions all estimated small *negative* coefficients for the distance to component mean. There were no significant regression results for either condition of the discrete whole-report task when data were separated by report order.

Finally, we repeated all analyses above with experimental data from the continuous task with orientation stimuli (Supplementary Fig. 13) and found no significant regressions with positive coefficients. Taken together, the many regression analyses we performed suggest that color reports were not consistently biased toward inferred ensemble mean in any condition of the whole-report task.

## Discussion

Here, we analyzed data from two whole-report visual working memory (VWM) experiments to characterize within-trial behavior and test the applicability of delayed estimation models. In contrast to models that assume independent encoding, we found that within-trial color reports were not independent. We also showed that this effect is either weaker or absent in task conditions using orientation stimuli, calling into question the assumption that color and orientation are equivalently encoded variables. Finally, we implemented two hierarchical Bayesian ensemble encoding models that explicitly include dependence between encoded stimuli and found no empirical evidence for the biases that they predict.

We began by replicating the continuous whole-report results reported by Adam, Vogel, and Awh, 2017 with data from our own discrete whole-report dataset. Although we successfully replicated several key results, our analyses of within-trial behavior in both datasets raise a potential issue with their conclusions. Specifically, Adam et al. found that late-trial report distributions were best described as uniform, and that participants typically self-reported that late-trial color reports were guesses. Reasoning that participants had no information about several items on each trial, they concluded that this was clear evidence for a fixed item limit in VWM. This has proven controversial, as some have argued that apparently uniform distributions could reflect very low-precision retrieval (Schneegans, Taylor, and Bays, 2020), and others have failed to reproduce late-trial uniform distributions altogether (Oberauer, 2022). Our results raise a different issue: we found that late-trial color reports were consistently biased away from early reports, which suggests that participants had *some* information about the colors they were reporting—even when self-reporting guesses.

Our analyses addressed the possibility that within-trial dependencies were indicative of hierarchical Bayesian ensemble encoding, but it is also possible that participants were using explicit or implicit task strategies that introduced dependency. For example, discrete task participants tended to consecutively report similar but not identical colors, suggesting that they were exploiting the absence of stimulus repeats within each trial. It has also been shown that certain whole-report stimuli can induce structured non-uniform guessing in late-trial reports (Ngiam et al., 2023), and we consider it probable that whole-report error distributions reflect both “memory” and “non-memory” processes. Unfortunately, post-hoc attribution of behavior to one process or another is not always straightforward—what if a participant intentionally guesses a color that they *do not* remember?—and distinguishing between behavior resulting from encoding structure versus task strategy raises challenges for the interpretation of VWM tasks that we will revisit below.

Adam, Vogel, and Awh, 2017 also reported very similar results for whole-report task conditions using color stimuli and those using orientation stimuli, and fit a set of models that assume a generic circular variable is encoded. Modeling color and orientation as equivalent is common (van den Berg et al., 2012; Schneegans, Taylor, and Bays, 2020), but recent work on has challenged the treatment of color as a circular variable (Schurgin, Wixted, and T. F. Brady, 2020). Although our results do not directly address the nature of color representations, the behavioral differences we identified between color and orientation conditions highlight the need to consider stimulus-dependent effects. It is also worth noting that there is considerable variation in visual working memory stimuli even when using the “same” stimulus type—few studies use standardized screen calibration methods for color (Bae, Olkkonen, et al., 2014), and orientation stimuli include Gabor patches ranging from −*π*/2 to *π*/2 (van den Berg et al., 2012) as well as various oriented “clock hands” and triangles ranging from −*π* to *π* (Adam, Vogel, and Awh, 2017; Bae and Luck, 2017; Utochkin and T. F. Brady, 2020).

Our focus on ensemble encoding as a potential cause of within-trial report dependencies was motivated by a rich body of VWM literature. The assumption of independent stimulus encoding has been challenged before using a variety of single-report working memory tasks (Jiang, Olson, and Chun, 2000; Kahana and Sekuler, 2002; Alvarez and Oliva, 2009; T. F. Brady and Alvarez, 2011; Orhan and Jacobs, 2013; Lew and Vul, 2015; Bae and Luck, 2017; Nassar, 2018) and at least one whole-report task (Utochkin and T. F. Brady, 2020), so it was unsurprising that we found evidence for dependence in the whole-report task. More surprising was that the hierarchical encoding models we implemented predicted report biases that were not present in either whole-report dataset. Our regression analyses of empirical reports failed to reproduce the consistent positive regression coefficents (report bias toward inferred ensemble or component mean values) that were present in HBM and BFMM simulations. In fact, a small number of regressions resulted in significant *negative* coefficents—this could be interpreted as bias away from ensemble or component means, but the effect was not consistent enough for us to draw such a conclusion. These results contrast with previous studies showing report biases toward mean stimulus size (T. F. Brady and Alvarez, 2011), mean horizontal position (Orhan and Jacobs, 2013), and mean 2-dimensional spatial position (Lew and Vul, 2015) that were accounted for by very similar hierarchical Bayesian encoding models. This discrepancy could possibly be explained by differences in stimulus type and task structure, but this interpretation is complicated by the results of Utochkin and T. F. Brady, 2020, where reports were biased toward mean orientation in a whole-report task.

It is important to note that ensemble encoding can take many forms, and that our results only contradict ensemble encoding models that assume a specific hierarchical Bayesian generative process. There are types of report bias—such as repulsion from dissimilar colors (Nassar, 2018)—that are not accounted for by hierarchical encoding but can still be considered evidence for ensemble encoding. Furthermore, a recent analysis of the same continuous whole-report dataset from Adam, Vogel, and Awh, 2017 found evidence for systematic biases that were explained by a non-hierarchical chunking model (Chunharas and T. Brady, 2023). Specifically, Chunharas and T. Brady, 2023 show that in trials identified by the model as chunkable, early color reports were biased toward the “gist” color and later color reports were biased away from the “gist” color. The “gist” in their analysis differs subtly from the ensemble and component means inferred by the hierarchical Bayesian models we implemented in that it involves the convolution of an empirically-derived psychophysical similarity function (Schurgin, Wixted, and T. F. Brady, 2020) with presented stimulus values. This method could be considered an empirical or phenomenological approach to identifying ensemble effects, in contrast to hierarchical Bayesian encoding models (T. F. Brady and Alvarez, 2011; Orhan and Jacobs, 2013; Lew and Vul, 2015) that make specific assumptions about the encoding structure used by participants. In our view, our results are compatible with those reported by Chunharas and T. Brady, 2023—although we did not find evidence for hierarchical encoding, the dependencies we identified are consistent with ensemble encoding more broadly, and we agree with their rejection of independent-item-based accounts of visual working memory.

## Conclusions

We analyzed data from two whole-report visual working memory tasks, and identified behavior that cannot be accounted for by models that assume independent encoding, equivalence between color and orientation stimuli, or hierarchical ensemble encoding. Given that these models were developed to fit single-report delayed estimation data and did not include sequential report mechanisms, it might be tempting to attribute such discrepancies to the idiosyncratic nature of the whole-report task. After all, the whole-report task plausibly suffers from the “task impurity” problem (Burgess, 1997) because it involves too many cognitive processes—attention, decision-making, strategies, etc—that are incidental to the structure of VWM encodings.

This problem is not unique to the current study, and raises a difficult dilemma. Should we reject more complex working memory tasks in the pursuit of “pure” working memory encoding models that may not generalize, or should we attempt to model the many processes responsible for idiosyncratic phenomena specific to the whole-report task? There is a growing recognition in the field that the proliferation of diverse VWM tasks, models, and empirical findings is outstripping our ability to develop theory capable of synthesizing them (Oberauer et al., 2018; Popov, 2023; Ngiam et al., 2023). In this context, neither option seems appropriate. Instead, we echo calls for a more integrative approach to theory development (Ngiam et al., 2023) and caution against inferring fundamental principles of visual working memory using data from any individual laboratory task.

## Additional acknowledgments

This preprint was created using the LaPreprint template (https://github.com/roaldarbol/lapreprint) by Mikkel Roald-Arbøl .

We would like to sincerely thank K Adam, E Vogel, and E Awh for sharing their experimental data publicly and therefore for making this project possible.

## Author contributions

Conceptualization: B.C., D.S., M.P., G.B.; Methodology: B.C., G.B.; Software and Analysis: B.C.; Writing - original draft: B.C.; Writing - review & editing: B.C., D.S., M.P., G.B.; Supervision: G.B., M.P.; Project administration: G.B.; Funding acquisition: G.B.

## Supplementary figures

**Figure 8.**
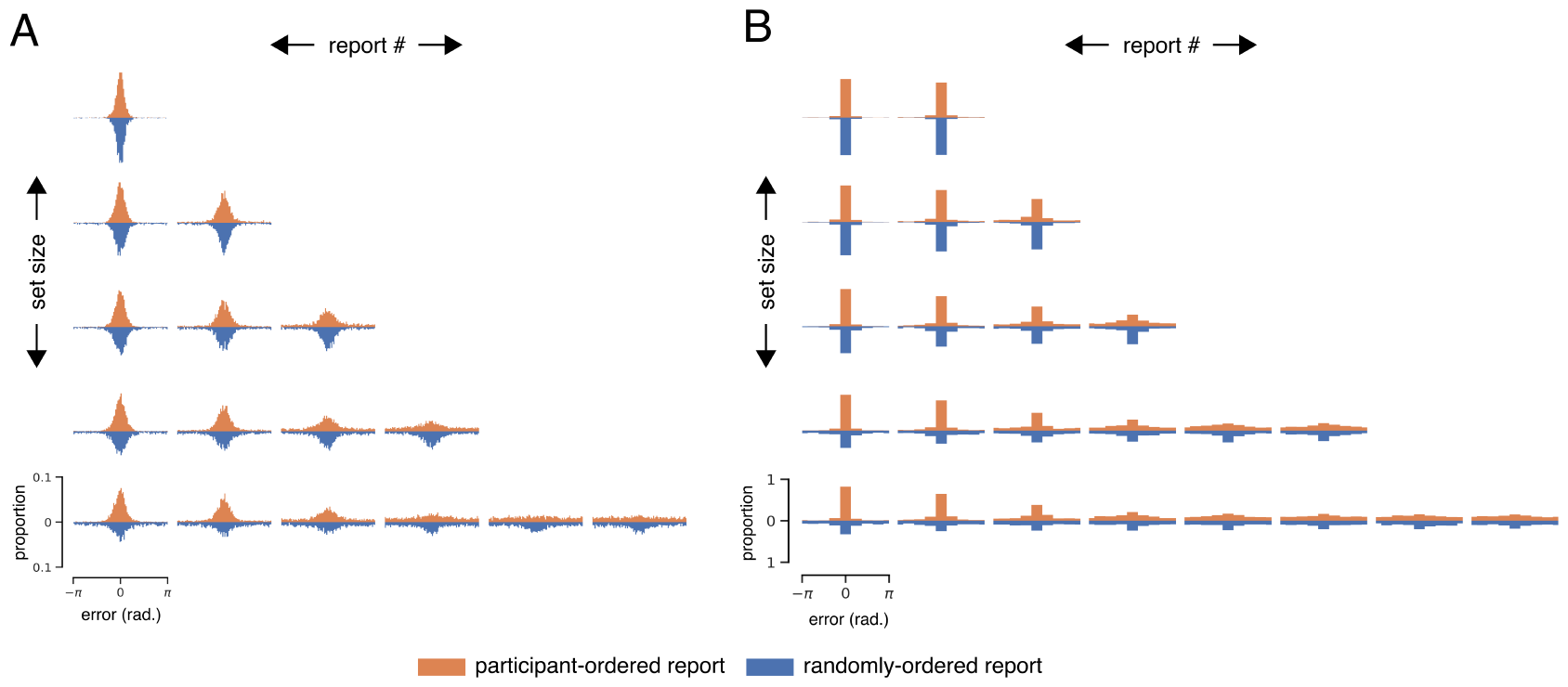
Color report error distributions for all set sizes, separated by report order. Includes all participants. **A**. Continuous error distributions. 360 possible error values divided into 90 bins for visualization. Orange and blue histograms show errors from participant-selected and randomly-generated report conditions, respectively. **B**. Discrete error distributions. Same as A, but bins correspond to all possible error values.

**Figure 9.**
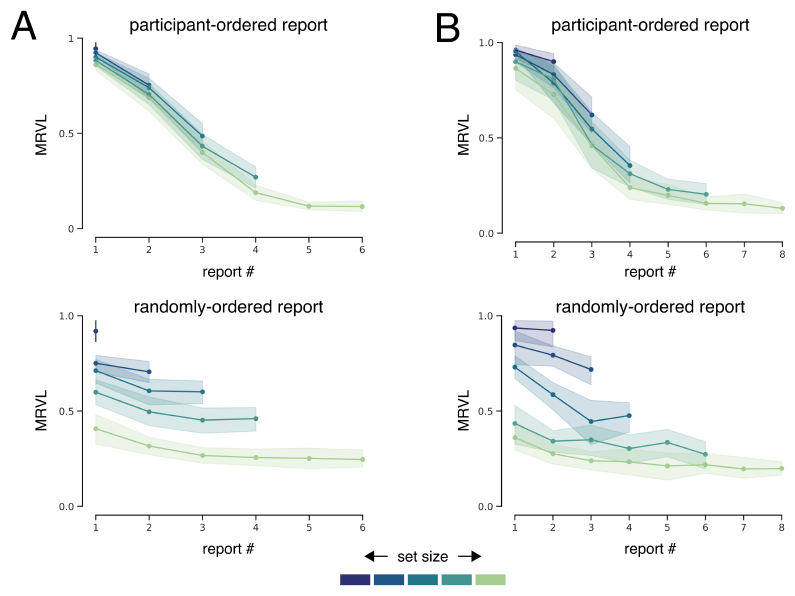
Mean resultant vector length (MRVL) of color report error distributions, separated by report order. **A**. Mean MRVL averaged across participants for the continuous whole report task. Shaded bars indicate +/- SEM. **B**. Same as A, but for the discrete task.

**Figure 10.**
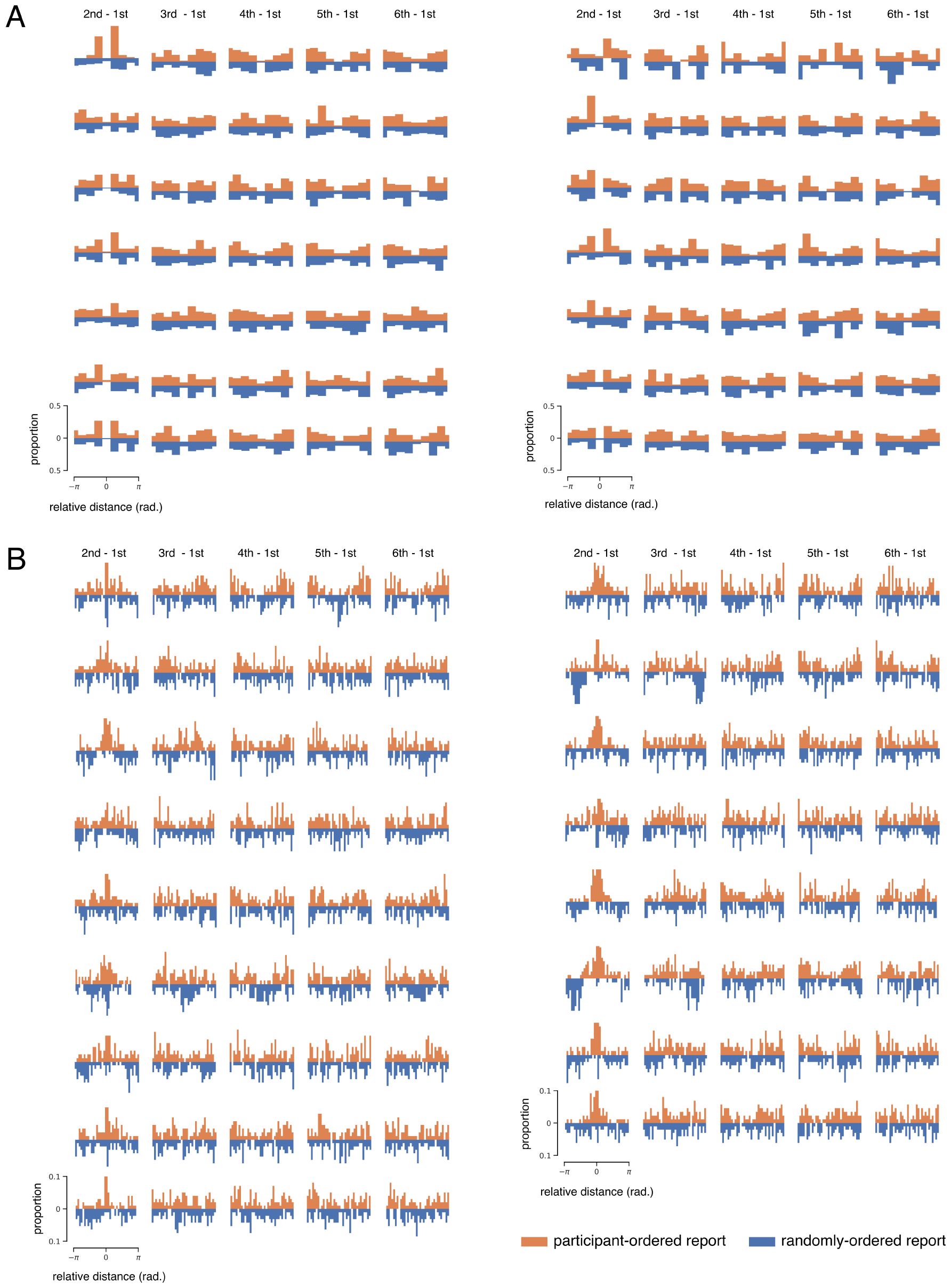
Within-trial joint color report distributions, separated by participant. **A**. Distribution of relative distances for the discrete task (set size 6). Each row shows data for one participant. **B**. Same as A, but for the continuous task.

**Figure 11.**
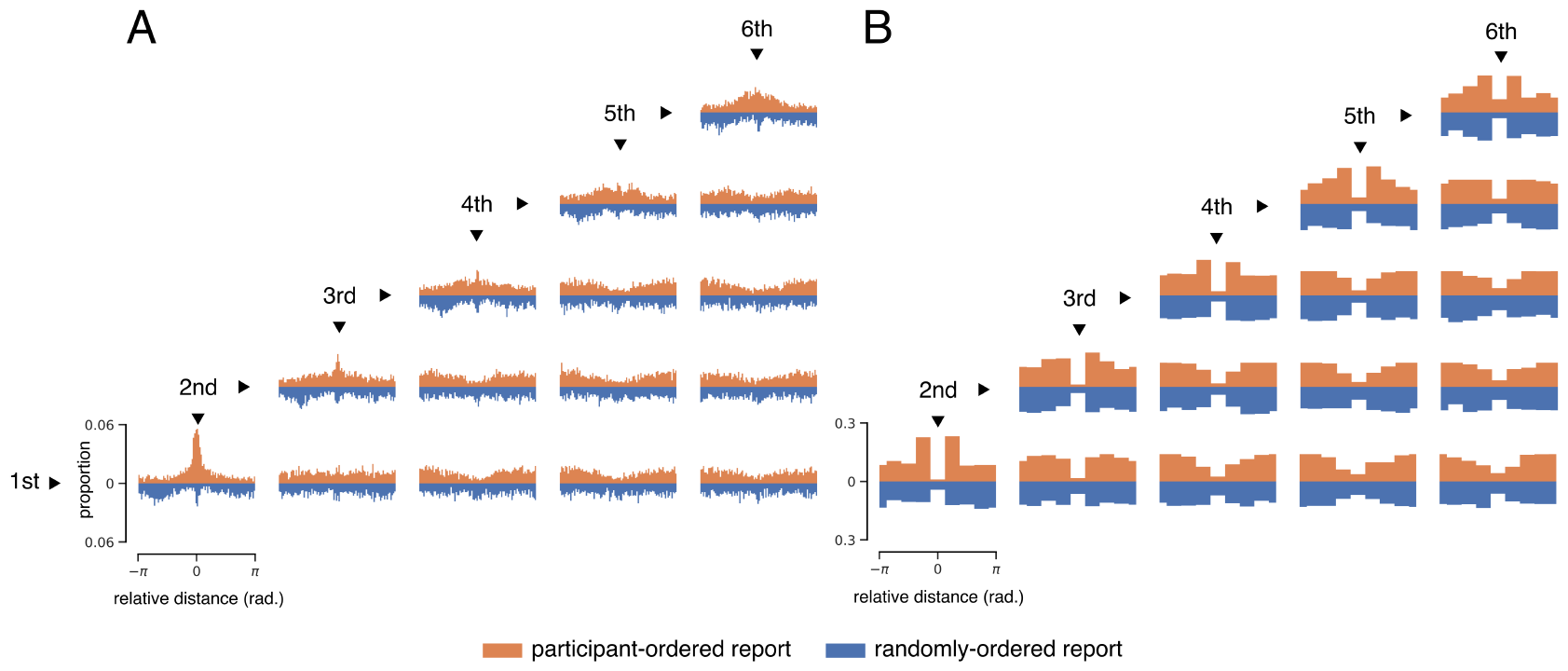
Within-trial joint color report distributions for all set sizes, collapsed across participants. **A**. Distribution of relative distances for the continuous task. Each row shows data for one set size. **B**. Same as A, but for the discrete task.

**Figure 12.**
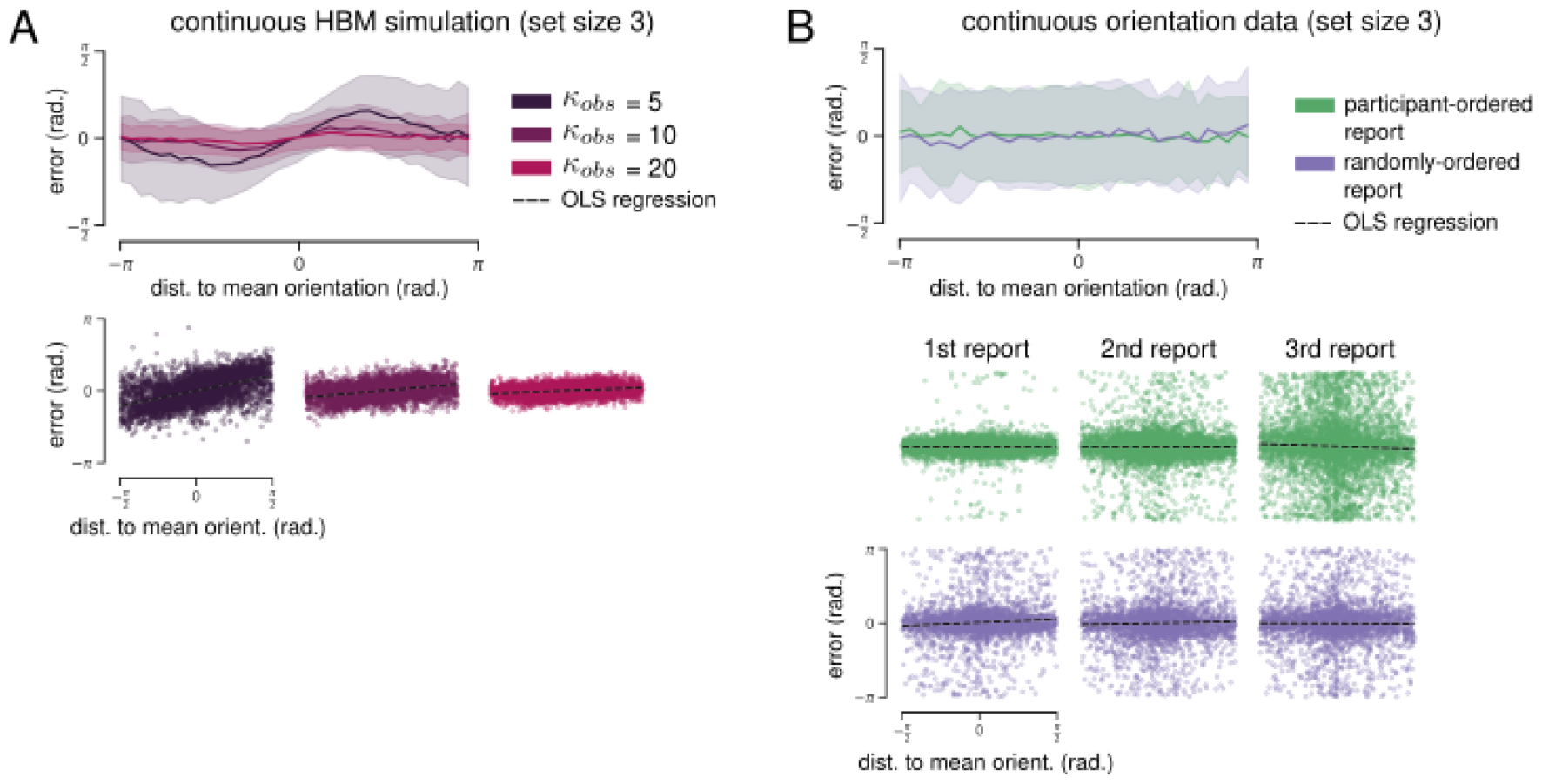
Hierarchical Bayesian model (HBM) simulation results for continuous task with orientation stimuli. **A**. *Upper:* Biases predicted by HBM simulations of the continuous task. Mean report error plotted as a function of the reported orientation’s distance to the mean orientation. 360 error values are divided into 90 bins for visualization, and shaded area shows standard deviation. *Lower:* OLS regression of simulated report error on distance to mean orientation (dashed black lines). **B**. *Upper:* Empirical biases for the continuous task at set size 3 (collapsed across all reports). *Lower:* OLS regression of empirical report error on distance to mean orientation. Each column shows results for a different report number.

**Figure 13.**
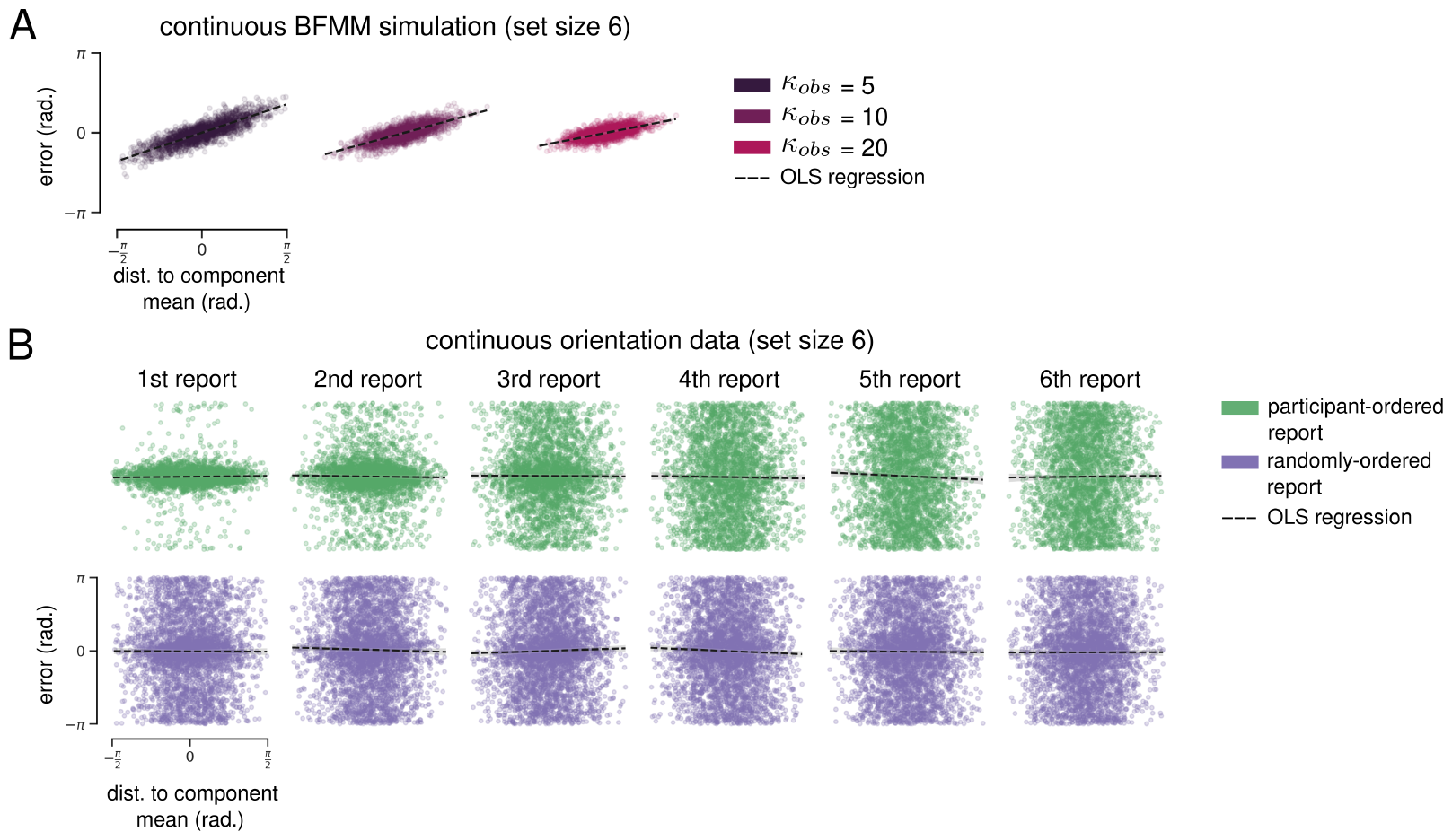
Bayesian finite mixture model (BFMM) simulation results for continuous task with orientation stimuli. **A**. *Upper:* Biases predicted by BFMM simulations of the continuous task. Mean report error plotted as a function of the reported orientation’s distance to the component mean. 360 error values are divided into 90 bins for visualization, and shaded area shows standard deviation. *Lower:* OLS regression of simulated report error on distance to component mean (dashed black lines). **B**. OLS regression of empirical report error on distance to component mean. Each column shows results for a different report number.

## Footnotes

1 Note that the joint distribution of two circular variables lies on the surface of a torus. This means that a point in the upper-left or bottom-right of panels in Fig. 2B represents two values very near each other in circular space.

2 Note that baseline KL divergence estimates are higher for the color distributions—this is because participants in the orientation task completed 200 trials per set size (vs 99 for the color task).

3 Note that reports are not simulated sequentially—neither the HBM or the BFMM include a temporal or sequential component. For visualization of both simulations and data, the bias functions in Fig. 5 include all reports, but the simulation regression plots in Fig. 5A and C include only the “first” simulated report to facilitate comparison with data regressions, which are separated by report order.

4 Orhan and Jacobs, 2013 also implemented a non-parametric model that allows for infinite groups. The BFMM is a constrained version, but is equivalent when *K* is equal to the set size.

